# Vegetative induction increases plant resistance to antagonistic insect frugivores

**DOI:** 10.1101/2024.12.27.628428

**Authors:** Deidra J. Jacobsen, Carmen D. Hewko

## Abstract

Plant fitness is shaped by interactions with insect mutualists and antagonists. Vegetative (leaf) herbivory often results in increased allocation of resources to defense in order to deter further damage. This allocation to defense can reduce floral or reproductive allocation. Reductions in floral allocation can have negative effects on pollinator attraction and therefore decrease plant fitness. However, defense-induced changes in reproductive tissues may also be protective against antagonistic fruit-feeding insects (frugivores). This may have important implications for plant fitness when plants experience multiple types of insect damage, but the potential protective effect of herbivory against frugivory is not well-understood.

In this study, we tested the prediction that herbivory mediates interactions between plants and their antagonistic frugivores. In the greenhouse, we manipulated vegetative induction in *Physalis pubescens* (Solanaceae) and measured the effects on plant fitness via changes in floral and fruit allocation, herbivore resistance in the next generation, and deterrence of an antagonistic frugivore (*Chloridea virescens* (Lepidoptera)).

Vegetative induction in *P. pubescens* enhanced plant resistance to *Chloridea virescens* frugivores with minimal negative effects on plant reproductive traits. Vegetative induction reduced *C. virescens* larval growth rate on fruits. *Chloridea virescens* also avoided induced plants for larval fruit feeding and oviposition. While leaf induction reduced flower size, induction did not negatively affect fruit size, seed set, or seed germination. Furthermore, offspring of induced plants showed increased resistance to *Manduca sexta* leaf herbivory. These findings indicate that herbivore-induced resistance may benefit plant fitness when plants are under simultaneous pressure from herbivores and frugivores.

## Introduction

Although herbivory often negatively affects plant reproduction, the consequences of leaf feeding and defensive induction on plant fitness are context-dependent and shaped by interactions with both mutualists and antagonists (Coley et al. 1985, Johnson et al. 2015). Plants must balance pollinator attraction and herbivore avoidance, which can result in conflicting selection on plant reproductive and defensive traits (Jacobsen and Raguso 2018). Leaf herbivory is known to alter vegetative defensive traits that protect against further feeding, but this resistance can come at a reproductive cost of fewer or smaller flowers, which can decrease pollinator attraction (Lehtilä and Strauss 1999, Mothershead and Marquis 2000). Although the impact of herbivory on floral and reproductive resources is often examined from the pollinator perspective, plants interact with a variety of insects, including flower feeders (florivores), fruit feeders (frugivores), and seed feeders (granivores) (Herrera et al. 2002, Poelman and Dicke 2014, Nelson and Whitehead 2021). Interactions among insects sharing a host plant can affect plant fitness as well as fitness of the interacting insects. Therefore, understanding the role of herbivory in mediating interactions with other insects is crucial to determine the net outcome of herbivory on plant fitness.

Despite temporal and/or spatial separation on the same host plant, leaf-feeding herbivores can indirectly affect flower or fruit visitors though plant-mediated interactions. Induced plant resistance after herbivory is often coordinated by the production of jasmonates (e.g. methyl jasmonate and jasmonic acid (JA)) (Eckardt 2008). Vegetative induction may alter leaf, flower, and fruit production as well as secondary chemistry, nutritional quality, or attractiveness of these tissues to insects. These changes can reduce insect success feeding on defended plants through reductions in larval growth or deterrence of larval feeding. Plant defenses also inform insect oviposition choices to maximize offspring success (i.e. optimal foraging and optional oviposition) (Jaenike 1978, Cipollini and Levey 1997, Scheirs 2002, McArt et al. 2013, Whitehead and Bowers 2013). Whether insects are repelled by leaf induction depends on whether defenses are localized or systemic, whether leaf and reproductive secondary chemistry is correlated, and type of insect.

Herbivory often results in correlated increases in nectar or pollen defensive compounds that can negatively affect pollinator preference and performance, harming plant fitness (Adler et al. 2006, Kessler and Halitschke 2009, Stevenson et al. 2017, Chautá et al. 2017). Other herbivory-induced changes that affect insect success and behavior can benefit the plant by reducing insect damage (Strauss and Irwin 2004, McArt et al. 2013). For example, in *Brassica nigra*, leaf herbivory increased glucosinolates in floral tissues, which reduced florivore performance (Rusman et al. 2022). In *Oenothera biennis*, leaf herbivory by an invasive beetle reduced the amount of damage from larvae of two moths that feed on flower buds (McArt et al. 2013). Likewise, changes in plant chemistry or reproductive allocation can affect frugivore attraction and behavior, but whether this is beneficial or detrimental to the plant depends on whether frugivores disperse or destroy seeds.

While fruit consumption by seed-dispersing frugivores (e.g. birds) commonly serves a beneficial role for plants, insect fruit feeding is often antagonistic, reducing plant fitness through loss of fruits that are costly for the plant to produce (Herrera 1982, Andersen 1988, Sallabanks and Courtney 1992). If induction at the vegetative stage increases plant resistance to insects that feed on fruits, this protection could outweigh the negative impact of herbivory on plant fitness in certain contexts. Reductions in antagonistic frugivory after herbivore induction would be particularly beneficial for plants with high tolerance to herbivory or that experience high frugivore pressures. Thus, it is necessary to determine whether vegetative defensive induction can decrease frugivore fitness without a large negative effect on plant fitness.

Herbivory can have additional impacts on plant fitness that extend beyond deterrence/harm of frugivores. Herbivore-induced changes in fruit size or chemistry may impact plant fitness in the next generation though changes in seed size, germination success, seedling defense, or seed longevity/resistance to pathogens (Strauss et al. 2002, Whitehead and Bowers 2013, Hernandez-Cumplido et al. 2016, Dalling et al. 2020, Whitehead et al. 2022). Herbivory can also change the timing of fruit and seed set (Redman et al. 2001, Whitehead and Poveda 2011). Understanding the fitness impacts of herbivory requires quantifying how herbivory affects both plant reproductive allocation and interactions with antagonistic insects (including herbivores and frugivores).

In this manipulative greenhouse study, *Physalis pubescens* (Solanaceae) were induced at the vegetative stage to measure the effects on plant reproduction and preference and performance of the frugivore *Chloridea virescens* Fabricus (Lepidoptera: Noctuidae). Leaf induction at the vegetative stage reduced flower size without negatively affecting fruit size, seed set, or seed germination. Importantly, vegetative induction deterred frugivore oviposition and larval feeding and reduced growth of frugivore larvae. Offspring of induced plants also had higher resistance to leaf herbivory by *Manduca sexta* L. (Lepidoptera: Sphingidae), a specialist leaf herbivore. These findings provide important evidence that herbivore induction can enhance plant resistance to antagonistic frugivores and increase resistance to herbivores in the next generation, thereby offsetting the potential negative effects of herbivory on pollinator attraction and growth.

## Materials and Methods

### System

*Physalis pubescens* L. is an annual, self-compatible plant that is host to a variety of leaf and fruit feeding insects, including the herbivore *Manduca sexta,* a specialist on Solanaceous species (Madden 1945, Menzel 1951, Waterfall 1967). *Manduca sexta* larvae have reduced growth rates when feeding on leaves of induced plants of many *Physalis* species, including *P. pubescens (Jacobsen 2022)*. *Physalis pubescens* produces constitutive and inducible secondary compounds that protect against insect herbivores (Doan et al. 2004, Zhang and Tong 2016).

*Physalis* fruits are attacked by a specialist frugivore, *Chloridea subflexa* Guenée (formerly genus *Heliothis*) (Pogue 2013)*. Chloridea subflexa* larvae use the papery fruit husk for protection from enemies while feeding on the fruit (Oppenheim and Gould 2002). The related generalist frugivore *Chloridea virescens* also feeds on *Physalis* and is a broader-scale crop pest (Sitchawat and Richard 1980). Many *Physalis* species are grown agriculturally and fruit damage from *Chloridea* larvae is economically harmful (Campos De Melo et al. 2017). *Chloridea virescens* has been used for feeding and oviposition studies with *Physalis* and is used here because it is easily reared in the lab and commercially available (Tingle et al. 1990, Sheck and Gould 1993).

### Plant culture

*Physalis pubescens* seeds were soaked in gibberellic acid (1g/1L) for two days before moving to flats (cell size 3.8 x 5.7 cm) filled with Redi-earth germination mix (Sun Gro Horticulture, Agawam, MA, USA). Two seed sources were used: accession 64400897GI from the Germplasm Resources Information Network (GRIN) originally from MN, USA and mixed seeds from multiple fruits from Orangeburg County, SC, USA. Flats were bottom watered in the greenhouse at the University of Utah (Salt Lake City, UT, USA). Approximately four to six weeks after germination, juvenile (pre-flowering) plants were moved to 3.8-liter pots filled with Lambert LM-3 all-purpose mix (Lambert Peat Moss, Québec, CA). Plants were provided with supplemental lighting to maintain 16:8 light:dark conditions. After one to two weeks of acclimation in the larger pots, leaf number was counted and plants within each seed source were randomly assigned to either the control (constitutive resistance) or induced resistance treatments (N = 118 plants total). Plant locations were randomized weekly to avoid positional effects. These plants were used for measurements of plant size/reproduction and *C. virescens* larval and oviposition assays. Seeds from these plants were collected for the remaining assays of seed germination and transgenerational resistance to *M. sexta* as described below.

### Plant induction treatments

Jasmonic acid (JA) was used to induce plant resistance in a standardized way. Exogenous jasmonates reliably induce resistance in Solanaceous species, including *Physalis* (Thaler et al. 1996, Doan et al. 2004, Jacobsen 2022). Plants in the induced treatment were treated with a combination of JA and mechanical tissue removal to mimic both the chemical and physical effects of herbivory. Plants in the induced treatment were sprayed with 1 mM JA in dH_2_O to run-off (Cayman Chemical, Ann Arbor, Michigan) and eight holes were punched in the leaves (two holes per leaf on four leaves per plant). Plants in the control treatment were sprayed with an equivalent amount of EtOH in dH_2_O to control for the solvent used to dissolve the JA. The JA and control spray treatments were repeated one week later.

### Plant size and reproductive measurements

When plants flowered (approximately two weeks after treatments), root crown diameter was measured as a proxy for overall plant size. Flower size (corolla width) was measured for the first three flowers and these flowers were tagged for measurements at the fruit stage. To compare total reproductive allocation, total flower and fruit numbers were tallied for all plants.

To quantify seed number and size, the first three fruits from each plant were weighed and dissected to remove seeds. A cohort of ten filled seeds from each of the first three fruits on each plant was weighted. To determine whether induction affected seed germination success or time to germination, germination assays were conducted at The Conservatory at Miami University in Hamilton, OH, USA (hereafter: The Conservatory). A cohort of ten seeds from one fruit per plant was placed in lidded two-ounce cups filled with distilled water and placed in the greenhouse under ambient light conditions during February-April with supplemental heat to keep temperatures above 13 °C at night. Time to seed germination was recorded for all cups at 23 timepoints over six weeks (after which no additional seeds germinated). Percent germination and days until germination of the first seed and last seeds was recorded.

### Measurements of transgenerational plant resistance

Because maternal induction can affect offspring fitness though not only seed size and provisioning but also through transgenerational defense, we tested whether offspring of induced maternal plants differed from uninduced plants in their resistance to the leaf herbivore *M. sexta*. The seeds for these assays came from fruits generated on induced or control maternal plants during the induction treatments at the University of Utah. These seeds were germinated at the The Conservatory in the same manner as the germination assays. Once seedlings had cotyledons, they were moved to flats (10 cm wide x 8 cm high, Foxany, Guangdong CN) filled with Pro-Mix Bx growing medium (Premier Tech Horticulture, Quakertown, PA USA) and bottom watered under ambient light conditions and supplemental heat to keep nighttime temperatures above 13 °C. Once they reached the juvenile stage, plants were transplanted to 2-liter pots and watered daily. Plants were grown for approximately three weeks until they reached the early budding/flowering stage.

For these trials, *M. sexta* larval growth rate was used as a proxy for plant resistance, with high larval growth rates indicating weak resistance and low larval growth rates indicating higher plant resistance (Haak et al. 2014, Jacobsen 2022). *Manduca sexta* larvae used for the feeding trials came from a colony at Miami University originally from Carolina Biological Supply (Burlington, NC USA) and were reared individually at 15:9 light:dark conditions at 22 °C on artificial diet until the second instar stage when they were used for the trials (tobacco hornworm diet F9783B, Frontier Agricultural Sciences, Newark, DE USA). Three hours prior to feeding trials, larvae were removed from diet to clear gut contents and increase hunger before weighing. Larvae were added individually to lidded two-ounce cups with moist filter paper and an upside-down cut leaf from either the maternal control or maternal induction plants. Larvae fed for 24 hours before taking final larval mass to calculate larval growth rate [log(final mass/initial mass)]. Larval growth rates were averaged across two larvae per plant and within plants from the same maternal family (N = 60 larvae; 17 unique maternal plants per treatment and 1-4 plants per maternal family). Larvae that did not eat or lost weight were removed from the trials (larvae do not feed during molting) (Jacobsen 2022).

### Frugivore larval growth rate based on prior plant induction

To determine whether plant vegetative induction affects resistance to frugivores, growth rates of *Chloridea virescens* larvae on fruits from control or induced plants were compared. Fruits for these feeding trials were autonomously selfed on the control and induced plants used for the plant size and reproductive measurements. Fruits were harvested at the preferred stage for *Chloridea* larval feeding (swollen fruits with immature seeds) (Campos De Melo et al. 2017). For each plant (N = 38 per treatment), three fruits were weighed and then placed on moist filter paper in separated lidded two-ounce containers. The *C. virescens* were reared in an environmental chamber at The University of Utah at 22 °C on a 14:10 light:dark cycle. Larvae were fed a pinto bean artificial diet (modified from Shorey and Hale 1965) prior to trials. Six-day old larvae were starved for two hours prior to the feeding trials to clear gut contents and increase hunger (Appendix S1: Table 1). After three days of feeding on the fruits, larvae were removed and starved for two hours to clear gut contents before recording final larval mass. Larvae that did not consume any of the fruit or died during the trial were excluded from calculations of growth rates, but lack of feeding was recorded to calculate fruit acceptance. Growth rate for each larva was calculated as log(final mass/initial mass) and the three larval growth rates on fruits from the same plant were averaged (Haak et al. 2014).

The three-day time period for these feeding assays was chosen to minimize the effect of any potential short-term compensatory strategies that could obscure the negative effects of feeding on induced fruits (Behmer 2009). Since larvae may alter their consumption based on fruit quality or defense, the percent change in fruit mass was compared between control and induced treatments to estimate the amount of the fruit eaten by the frugivore. Fruits were measured before and after the feeding trial to calculate proportion change in fruit mass ((final fruit mass – initial fruit mass)/(initial fruit mass).

*Chloridea* eggs are often laid on leaf tissue in addition to reproductive tissue (Benda et al. 2011), so an additional set of *C. virescens* feeding trials was done on control and induced leaf tissue to verify that the frugivores do not also act as leaf herbivores. In this feeding trial, larvae initiated feeding on leaves in the absence of fruits but the majority of larvae failed to gain weight feeding on leaves, especially when feeding on induced leaves (Appendix S2).

### Larval fruit choice assays

Because *Chloridea* larvae need to consume multiple fruits to pupate, fruit choice assays were used to determine whether leaf induction affects frugivore feeding choices (Benda et al. 2009). One week old *C. virescens* larvae were starved for two hours then placed in the center of a lidded four-ounce cup. Two size-matched fruits in their husks (one control and once induced) were placed across from each other in the cup (N = 100 larvae tested on 89 unique combinations of fruits from control and induced plants). Fruits were checked for feeding at 24 and 48 hours. After 48 hours, when the majority of larvae had chosen a fruit, fruit damage and active larval feeding were used to quantify *C. virescens* fruit choice. Because larvae often chew a hole through the husk to enter a fruit for feeding, husks were inspected for holes to determine whether larval husk sampling correlated with fruit choice.

### Adult frugivore oviposition choice

Maternal oviposition choice largely defines neonate feeding location (Benda et al. 2011). Therefore, 24-hour binary choice assays were conducted to quantify whether mated female *C. virescens* moths preferred to lay eggs on control or induced plants. Two-to-three-day old adult moths were mated in 3.8 liter containers overnight with access to a nine percent sucrose solution. Following mating, females were placed in individual cages in the greenhouse (34x34x61 cm, BioQuip Products 1466BV, Rancho Dominquez, CA, USA) with size-matched control and an induced plants on opposite sides of the cage. Plants had flowers and fruit (N = 17 pairs of plants). After 24 hours, moths were removed to count the number of eggs on each plant. If no eggs were laid, the moth was replaced with another moth to get data for all plant pairs.

### Statistical analysis

All analyses were done using R v. 4.3.2 (R Core Team 2013). Generalized linear models (GLM) and linear models (LM) were run in the stats package. Shapiro tests (shapiro.test) and Levene’s tests (leveneTest, “car” package) were used to check normality and homogeneity of variance (Fox, J. and Weisberg, S. 2019). Non-normal data was log-transformed. The describeBy function in the “psych” package was used to compare groups (Revelle 2024). The dispersiontest function in the “AER” package was used to check for overdispersion in the Poisson regressions (Kleiber and Zeileis 2008). If overdispersed, the quasi-Poisson distribution was used.

Because plant reproductive output is often positively correlated with plant size, root crown diameter was included as a fixed effect in plant reproductive trait models. Seed source was included as a fixed effect in plant trait models to account for any potential variability between the two accessions used. As described below, seed source did not significantly impact measurements of plant size, flower number or size, fruit size, seed number or size, or germination. Therefore, seed source was not included in insect growth and choice analyses.

Linear models were used to test whether root crown diameter (log-transformed) differed based on leaf induction treatment. A Poisson GLM was used to analyze whether vegetative induction altered flowering time. Separate quasi-Poisson GLMs were used to test if total flower or fruit number differed between control and induced treatments. To determine if there were differences in flower or fruit size, separate linear models were run for flower width and log-transformed fruit mass.

A quasi-Poisson GLM was used to test for differences in seed production between control and induced treatments for the first three fruits. Seed size in control and induced treatments was compared using linear models. Binomial GLMs were used to test whether the total proportion of seeds germinating depended on leaf induction treatment. The number of days to first and final seed germination for seeds from control or induced maternal plants was analyzed using separate quasi-Poisson GLMs.

Linear models were used to test if *C. virescens* larval growth rate differed when feeding on fruits from control versus induced plants. The proportional change in fruit mass was tested using a binomial GLM to determine whether larvae consumed different amounts of fruits from control or induced plants. Because differences in larval acceptance of fruits in either treatment could affect growth rate in the no-choice assays, a chi-squared test analysis (chisq.test()) was used to compare the number of larvae that did not eat the fruit offered in each treatment. Separate chi-squared analyses were used to compare the number of larvae choosing control fruits versus induced fruits for feeding and to determine whether fruit choice depended on whether or not the larvae sampled the husks. A paired, one-sided Wilcoxon test with continuity correction (wilcox.test()) was used to test if the number of fertilized eggs laid in the binary oviposition choice assays was significantly higher on control plants than on induced plants.

To assess whether maternal vegetative induction affects resistance to herbivores in the next generation, linear models were used to compare *M. sexta* larval growth rate on leaves from maternal control plant or induced plants.

## Results

### Effect of vegetative induction on plant growth and reproductive allocation

Plant induction with jasmonic acid at the vegetative stage (simulated leaf herbivory) reduced plant size at flowering. Root crown diameter at flowering was smaller for induced plants compared with control plants (mean diameter: control = 8.1 mm, induced = 7.1 mm) (LM: induction t = -2.568, P = 0.0121, seed source t = -1.683, P = 0.0964).

Vegetative induction delayed flowering but did not reduce flower or fruit number. Plants in the induction treatment flowered later than control plants (mean days between treatment and flowering: control =12.71 days, induced 14.33 days) (Poisson GLM: induction z = 1.992, P = 0.0463; seed source z = -0.604, P = 0.5460; root crown z = 0.626, P = 0.5316). Smaller plants produced fewer flowers than large plants, but there was no effect of induction on flower number (mean flower number: control = 127.4, induced =99.8) (quasi-Poisson GLM: induction t = 0.835, P = 0.406; root crown t=12.901, P < 0.0001, seed source t = -1.516, P = 0.134). Likewise, smaller plants produced fewer fruits, but vegetative induction did not significantly affect fruit number (mean fruit number: control 57.2, induced 48.1) (quasi-Poisson GLM: induction t= 0.508, P = 0.6128; root crown t = 7.887, P < 0.0001, seed source t = -2.311, P = 0.0238).

Vegetative induction reduced flower size but increased fruit size. Induced plants produced smaller flowers than control plants (mean flower width: control = 17.1mm, induced 15.73 mm) (LM: induction t = -3.632, P = 0.0005; root crown t = 0.357, P = 0.7223, seed source t = 1.197, P = 0.235). Induced plants and larger plants produced larger fruits (mean fruit size: control 0.58 g, induced 0.66 g) (LM: induction t = 2.065, P = 0.0425; root crown t = 4,187, P < 0.0001, seed source t = 1.489, P = 0.1407) (Figure 1).

**Figure 1.**
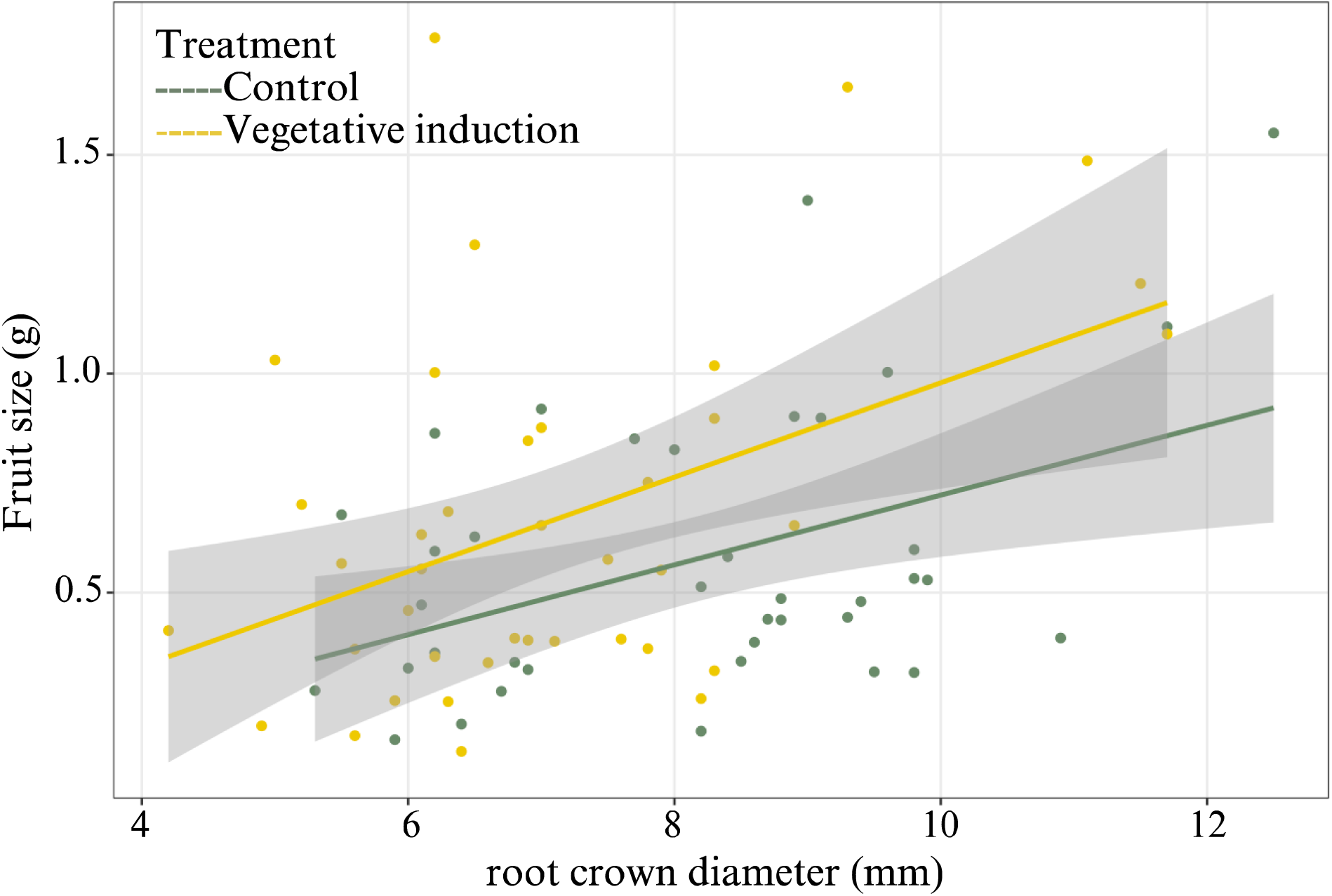
Fruits from *Physalis pubescens* plants in the vegetative induction treatment were larger than fruits of control plants (LM: induction t = 2.065, P = 0.0425; root crown t = 4,187, P < 0.0001, seed source t = 1.489, P = 0.1407). Lines display the linear regression for each treatment group and gray shading indicates the 95% confidence interval.

Fruits from induced plants contained more seeds per fruit than control plants, but seed size did not differ between treatments. Induced plants and larger plants produced more seeds per fruit (mean seed number per fruit: control = 67.34, induced = 86.3) (quasi-Poisson GLM: induction t = 2.090, P = 0.0401; root crown t= 2.069, P= 0.0421; seed source t = 1.623, P = 0.1090). Seeds from fruits on induced plants were similar in size to seeds from control plants (mean seed weight: control = 0.528 mg, induced = 0.481 mg) (LM: induction t = -1.546, P = 0.127, root crown t = 0.941, P = 0.350, seed source t = -0.956, P = 0.342).

### Effects of maternal vegetative induction on offspring seed germination and resistance

Plant induction did not change the proportion of seeds germinating or timing of seed germination. Seed germination success did not differ for seeds produced by maternal plants in the control versus induced treatments (mean proportion germinated: control = 0.64, induced = 0.58) (binomial GLM: induction z = -0.240, P = 0.810; root crown z = 0.046, P = 0.964, seed source z = 0.038, P = 0.970). The number of days until germination of the first seed did not differ based on maternal control or induction treatment (mean number of days to first seed germination: control = 12.3, induced = 13.4) (Poisson GLM: induction z = 1.423, P = 0.1547, root crown z = 1.885, P = 0.0594, seed source z = 1.493, P = 0.1356). Similarly, time until the last germination event did not differ based on maternal control or induction treatment (mean number of days to final seed germination: control = 22.32, induced = 23.94) (quasi-Poisson GLM: induction t = 0.349, P = 0.729, root crown t = -0.015, P = 0.988, seed source t =0.145, P = 0.886).

Maternal vegetative induction increased offspring resistance to leaf herbivory. *Manduca sexta* larval growth rates were lower when feeding on leaves if the maternal plant had been induced at the vegetative stage (LM: induction t=-2.365, P = 0.030) (Figure 2).

**Figure 2.**
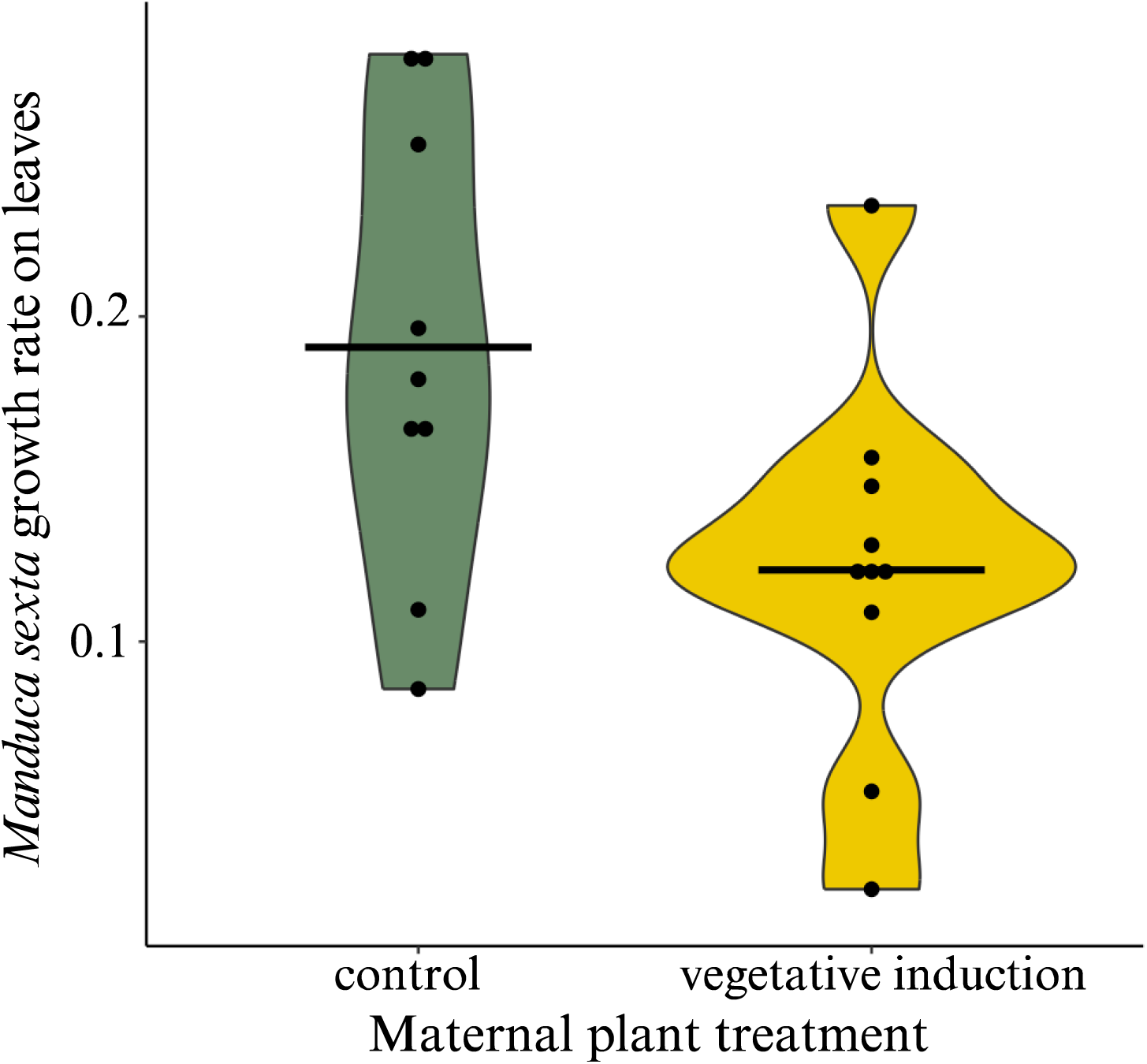
Offspring of maternally induced *Physalis pubescens* had higher leaf resistance to the herbivore *Manduca sexta* in a 24-hour no-choice feeding trial. Mean larval growth rate was calculated as log(final mass/initial mass) and averaged across two larvae per plant for one to four plants per maternal family. Dots represent growth rate per day on leaves from control plants (green fill) or maternally-induced plants (gold fill) and the line represents the treatment mean.

### Effect of vegetative induction on frugivore growth and behavior

Frugivore growth rates in the no-choice assay were lower for *C. virescens* larvae feeding on fruits from induced plants compared with control plants (mean growth rate over three days: control = 0.29, induced 0.19; LM induction: t = -3.917, P = 0.0002) (Figure 3). The amount of fruit consumed between treatments did not differ based on whether fruits came from control or induced plants (binomial GLM treatment z = -0.101, P = 0.9199). Larval *C. virescens* fruit acceptance during the no-choice fruit feeding assays did not differ for fruits from control or induced plants (N = 38 plants per treatment with 3 larvae tested per plant; control N= 8 did not eat, induced N= 5 did not eat) (X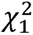=0.3712, P = 0.5424).

**Figure 3.**
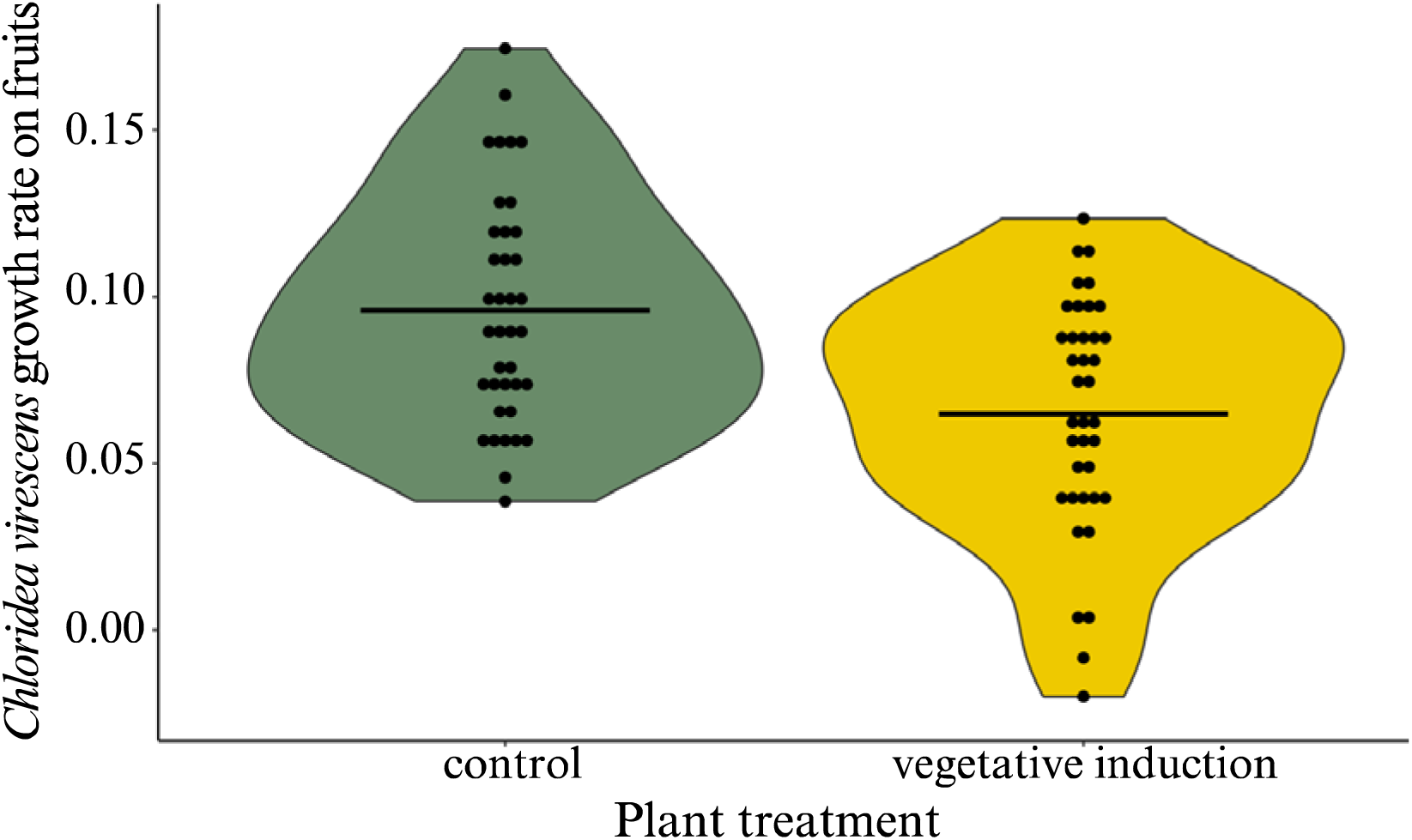
Violin plot showing a reduction in growth rate of *Chloridea virescens* larvae feeding on fruits from induced *Physalis pubescens* plants compared with those feeding on control fruits in a no-choice assay. Growth rates (log (final/initial mass) over a three-day period were averaged to calculate daily larval growth rate for a plant. Dots represent growth rate per day on fruits from control (green fill) or induced plants (gold fill) and the line represents the treatment mean.

In the binary choice assays*, C. virescens* larvae avoided feeding on fruits from induced plants. Larvae chose to feed on the induced fruit half as often as they chose the control fruit in the binary choice assays (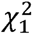=10, P = 0.0016). Husk sampling did not significantly alter fruit choice, as larvae that sampled both husks did not differ in their fruit choice from larvae that only sampled the husk of the fruit they chose (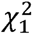=0.2756, P = 0.5996) (Figure 4). Only one larva moved to feed on the other fruit prior to consuming the majority of the first fruit chosen.

**Figure 4.**
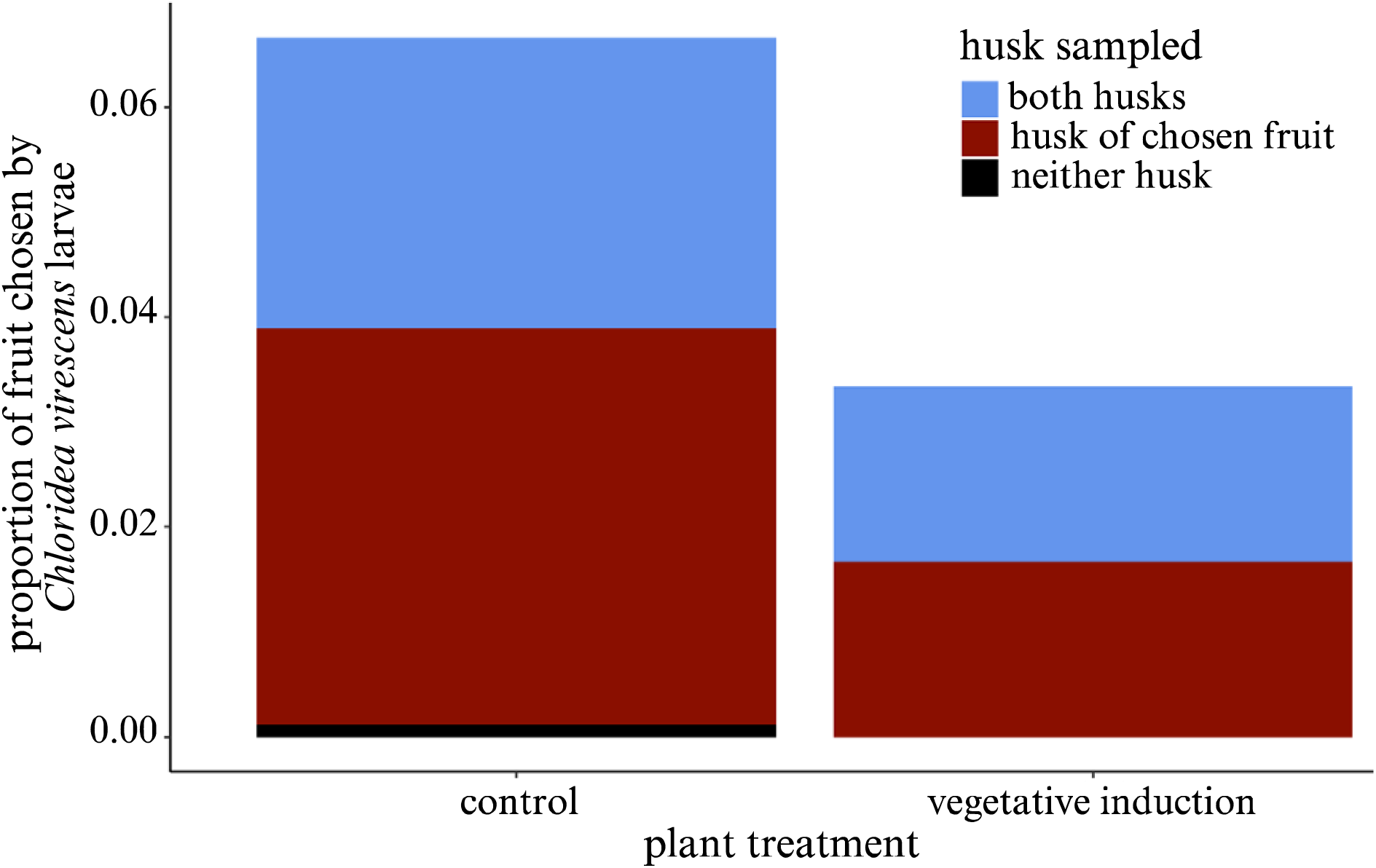
In 48-hour binary choice assays, *Chloridea virescens* larvae chose to feed on the fruit from the control plant over the fruit from the induced plant (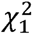=10, P = 0.0016). Of the N=100 larvae tested, two larvae died without making a fruit choice and eight larvae did not choose a fruit. Sampling of both husks did not significantly alter fruit choice; larvae sampled both husks (blue) or sampled only the husks of the fruit they chose (red) in similar proportions (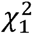=0.2756, P = 0.5996)

Plant induction deterred *C. virescens* oviposition. In binary choice assays with control and induced plants, mated female moths laid fewer eggs on induced plants than on control plants (mean egg numbers: control = 15.06, induced = 4.94) (paired Wilcoxon signed rank test: V = 106.5, P = 0.0245).

## Discussion

Interactions between plant defense and reproduction, leaf herbivores, and antagonistic frugivores can shape plant and insect fitness and co-evolutionary interactions. Much of the research on reproductive allocation and insect visitation focuses on the impacts on beneficial pollinators and seed dispersers rather than antagonistic frugivory (McCall et al. 2006). Here, we demonstrate that leaf herbivory can increase plant resistance to frugivore damage, even when leaf induction occurs prior to fruit damage. This indirect, host-mediated interaction between herbivores and frugivores sharing the same host plant is likely to increase plant fitness when under pressure from both herbivores and frugivores. Notably, this increase in resistance to frugivores occurred without costs to seed set or offspring germination and resulted in higher offspring resistance to leaf damage.

The fitness benefit of reduced fruit damage following herbivore feeding and/or leaf induction may outweigh potential negative effects on growth or attractiveness to pollinators. In this study, plant induction at the vegetative stage reduced flower size, which is known to negatively affect pollinator attraction in many systems (Moreira et al. 2019). However, in self-compatible species, such as *Physalis pubescens*, these species can still produce seeds even if they are less attractive to pollinators. Although induced plants had delayed flowering and reduced flower size, induction did not reduce fruit size or fecundity of the selfed fruits produced in the greenhouse. Induced plants produced larger fruits with more seeds per fruit than control plants. Seeds from fruits on induced plants germinated well and plants grown from these seeds had higher resistance to leaf herbivores than plants grown from uninduced maternal plants. In studies in other systems, herbivory-induced reductions in seed size and protein content reduced germination success (Hernandez-Cumplido et al. 2016) but plant induction did not affect seed size or germination in the present study. In the absence of intrinsic reductions in seed size or germination success, the benefit of reduced frugivory following plant vegetative induction is likely to increase fitness in the face of frugivore pressures.

Fruits from induced plants were more resistant to *C. virescens* frugivores, as shown by the reduced frugivore growth rate on fruits from induced plants and avoidance of induced plants for larval feeding and oviposition. The adult oviposition preference for control plants aligned with higher larval growth rates on control plants, consistent with optimal oviposition theory (Jaenike 1978). Weighing fruits before and after feeding in the three day no-choice trials did not show a significant difference in fruit mass (a proxy for amount of fruit consumed) when larvae were feeding on control or induced fruits. The decrease in *C. virescens* growth rate without a reduction in consumption indicates that leaf induction likely increases fruit defensive compounds or reduces nutritional quality for frugivores. Correlations between leaf and fruit defensive compounds may be due to adaptive roles of fruit secondary compounds as well as physiological or pleiotropic factors that constrain production of secondary compounds in leaves without corresponding increases in fruit secondary compounds (Whitehead and Bowers 2013).

Interestingly, although *Chloridea* larvae generally chew a hole though the fruit husk to gain entry to the fruit, husk sampling did not appear to alter fruit choice (Figure 4). Husks provide physical protection to frugivores (Sisterson and Gould 1999, Oppenheim and Gould 2002), however they do not appear to provide chemical or nutritional cues prior that inform *C. virescens* feeding on the fruit. Because *C. virescens* are generalist herbivores, husk sampling or fruit defensive chemistry may be more important to the specialist *C. subflexa*, which uses the husk for protection in the field (Sisterson and Gould 1999, Oppenheim and Gould 2002, Sell et al. 2021).

In the field there may be additional effects of vegetative induction on herbivore and frugivore feeding or oviposition preferences. For example, in this study fruits were size matched for the choice assays, but overall fruit size was larger on induced plants and this size difference may affect herbivore fruit choice in the field. There may be additional effects on other insects not considered in this study (McArt et al. 2013). Additionally, changes in fruit chemistry or nutritional content following vegetative induction may also affect interactions with fruit dispersers (Whitehead et al. 2022). *Physalis* fruits are often wind and water dispersed but seeds may also be dispersed by birds that consume the fruits (Olson 1999, Li et al. 2019). Because of the potential fitness effects of interactions between co-occurring conspecific and heterospecific insects on a host plant (Strauss and Irwin 2004, McNutt and Underwood 2016), future work in this system aims to describe these interactions under field conditions.

## Acknowledgements

Nicole Benda and GRIN provided seeds. Neil Vickers provided *C. virescens* and lab members assisted with moth care at the University of Utah. Plant care was provided by Christopher Morrow (Utah), Savannah Ballweg, Adrian Bennett, Aidan Oglesbee, and Emily Dudaitis at The Conservatory. CH received funding from the Miami University Herbarium.

## Author Contributions

CH assisted with the design for the germination and transgenerational assays and collected the data for these parts of the study. DJ collected data for the other projects and analyzed the data. DJ prepared the manuscript and CH contributed to revisions.

