## Appendix 1 for "Vegetative induction increases plant resistance to antagonistic insect frugivores"

Appendix 1. Chloridea virescens fed on leaf or artificial diet were starved for 3 hours to determine the time to clear gut contents based on loss of mass and production of frass. Nine day old larvae were weighted at the start (time = 0) and each hour during the three hours of starvation. For the first hour, all produced frass and lost an average of 3.4% of their body mass. Over the second hour, moths lost an additional 3.6% body mass and some but not all produced frass (in smaller amounts). Over the third hour they only lost another 1.2% in body mass and most produced only 0-1 piece of frass. Overall, Chloridea virescens cleared the majority of their gut contents in 2 hours and lose a total of 7% of their body weight in that 2 hours. Therefore, 2 hours was used as the starvation time prior to feeding trials to maximize gut clearing while minimizing added starvation stress.

| moth ID | larval age | part fed on | pre-mass (g) before starvation | mass after 1 hour starvation (g) | frass produced during hour 0-1 | mass after 2 hours starvation (g) | frass produced during hour 1-2 (previous frass wiped out) | mass after 3 hour starvation (g) | frass produced during hour 2-3 (previous frass wiped out) | 0-1hr %change in body mass | 1-2hr %change in body mass | 2-3hr %change in body mass | 0-2hr %change in body mass |
|---|---|---|---|---|---|---|---|---|---|---|---|---|---|
| L1 | 9 days | leaves | 0.0131 | 0.0124 | yes (>3 pieces) | 0.0122 | 1 piece | 0.0122 | no | 5.3 | 1.6 | 0 | 6.9 |
| L2 | 9 days | leaves | 0.0116 | 0.0112 | yes (>3 pieces) | 0.0103 | 1 piece | 0.0103 | no | 3.4 | 8 | 0 | 11.2 |
| L3 | 9 days | leaves | 0.0157 | 0.0147 | yes (>3 pieces) | 0.0147 | no | 0.0144 | 1 piece | 6.4 | 0 | 2 | 6.4 |
| L4 | 9 days | leaves | 0.013 | 0.0129 | yes (>3 pieces) | 0.0124 | 4 pieces | 0.0122 | 1 piece | 0.8 | 3.9 | 1.6 | 4.6 |
| L5 | 9 days | leaves | 0.0153 | 0.0145 | yes + webbing | 0.0132 | 2 pieces + webbing | 0.013 | no | 5.2 | 9 | 1.5 | 13.7 |
| L6 | 9 days | leaves | 0.0091 | 0.0087 | yes (>3 pieces) | 0.0073 | no | 0.0071 | no | 4.4 | 16.1 | 2.7 | 19.8 |
| L7 | 9 days | leaves | 0.0159 | 0.0154 | yes (>3 pieces) | 0.0154 | no | 0.0152 | 1 piece | 3.1 | 0 | 1.3 | 3.1 |
| L8 | 9 days | leaves | 0.0099 | 0.0099 | yes (>3 pieces) | 0.0097 | 2 pieces | 0.0097 | no | 0 | 2 | 0 | 2 |
| D1 | 9 days | artificial diet | 0.0228 | 0.022 | yes (>3 pieces) | 0.0215 | 1 piece + silk | 0.0211 | no | 3.5 | 2.3 | 1.9 | 5.7 |
| D2 | 9 days | artificial diet | 0.0169 | 0.0165 | yes (>3 pieces) | 0.0165 | 3 pieces | 0.0163 | 2 pieces | 2.4 | 0 | 1.2 | 2.4 |
| D3 | 9 days | artificial diet | 0.0114 | 0.0114 | yes (1 piece) | 0.0112 | no | 0.011 | no | 0 | 1.8 | 1.8 | 1.8 |
| D4 | 9 days | artificial diet | 0.0169 | 0.0165 | yes (>3 pieces) | 0.0161 | 1 piece | 0.016 | no | 2.4 | 2.4 | 0.6 | 4.7 |
| D5 | 9 days | artificial diet | 0.0191 | 0.0185 | yes (>3 pieces) | 0.0175 | 1 piece | 0.0171 | 1 piece | 3.1 | 5.4 | 2.3 | 8.4 |
| D6 | 9 days | artificial diet | 0.0279 | 0.0262 | yes (>3 pieces) | 0.0253 | 3 pieces + webbing | 0.0253 | no | 6.1 | 3.4 | 0 | 9.3 |
| D7 | 9 days | artificial diet | 0.0281 | 0.0267 | yes (>3 pieces) | 0.0264 | 1 piece | 0.0259 | no | 5 | 1.1 | 1.9 | 6 |
| D8 | 9 days | artificial diet | 0.0227 | 0.0218 | yes (>3 pieces) | 0.0217 | no | 0.0216 | no | 4 | 0.5 | 0.5 | 4.4 |
