## Appendix 2 for "Vegetative induction increases plant resistance to antagonistic insect frugivores"

**Appendix 2.** *Chloridea virescens* larval growth rate on leaf tissue.

Larval feeding trials on control and induced leaf tissue were done in a similar manner as the fruit feeding trials (N = 28 total larvae, N = 7 plants per treatment and N = 2 larvae per plant). Larvae were starved for two hours prior to feeding trials, weighted, then fed on mature leaf tissue in individual lidded two-ounce containers with a damp piece of filter paper to prevent the leaf from drying out. Larvae were weighed at the conclusion of the three day feeding period and the growth rate was calculated as log(final/initial mass) and two larval growth rates per plant were averaged. These growth rates were analyzed using linear models (lm()) in R.

The mean growth rate per day on control leaves was 0.01 and the mean growth rate on induced leaves was significantly lower at -0.02 (lm: t = -3.612, P = 0.0034). Sixty-four percent of the larvae failed to gain weight or lost weight over the three-day feeding period.
